# Co-regulation of alternative splicing by hnRNPM and ESRP1 during EMT

**DOI:** 10.1101/301267

**Authors:** Samuel E. Harvey, Yilin Xu, Xiaodan Lin, Xin D. Gao, Yushan Qiu, Jaegyoon Ahn, Xinshu Xiao, Chonghui Cheng

## Abstract

The epithelial-mesenchymal transition (EMT) is a fundamental developmental process that is abnormally activated in cancer metastasis. Dynamic changes in alternative splicing occur during EMT. ESRP1 and hnRNPM are splicing regulators that promote an epithelial splicing program and a mesenchymal splicing program, respectively. The functional relationships between these splicing factors in the genome-scale remain elusive. Comparing alternative splicing targets of hnRNPM and ESRP1 revealed that they co-regulate a set of cassette exon events, with the majority showing discordant splicing regulation. hnRNPM discordantly regulated splicing events show a positive correlation with splicing during EMT while concordant splicing events do not, highlighting the antagonistic role of hnRNPM and ESRP1 during EMT. Motif enrichment analysis near co-regulated exons identifies guanine-uridine rich motifs downstream of hnRNPM-repressed and ESRP1-enhanced exons, supporting a model of competitive binding to these cis-elements to antagonize alternative splicing. The set of co-regulated exons are enriched in genes associated with cell-migration and cytoskeletal reorganization, which are pathways associated with EMT. Splicing levels of co-regulated exons are associated with breast cancer patient survival and correlate with gene sets involved in EMT and breast cancer subtypes. These data identify complex modes of interaction between hnRNPM and ESRP1 in regulation of splicing in disease-relevant contexts.

## INTRODUCTION

Alternative RNA splicing is a fundamental mechanism of functional genome diversity that enables the nearly 21,000 protein-coding genes in the human genome to give rise to over 100,000 transcripts (ENCODE Project Consortium 2012; Harrow et al. 2012). Deep transcriptome sequencing has revealed that approximately 95% of all human multi-exon transcripts can undergo alternative splicing, positioning alternative splicing as a critical form of post-transcriptional gene regulation in a variety of cellular and biological processes (Pan et al. 2008; Wang et al. 2008; Barash et al. 2010). Dysregulation of alternative splicing is increasingly implicated in a variety of human diseases, including cancer progression and survival (Liu and Cheng 2013; Cieply and Carstens 2015).

Alternative splicing has emerged as a central regulatory process during the epithelial-mesenchymal transition (EMT) (Warzecha et al. 2010; Brown et al. 2011; Shapiro et al. 2011; Reinke et al. 2012; Yang et al. 2016). EMT is a developmental program whereby epithelial cells transit to a mesenchymal phenotype, which occurs in natural processes such as organogenesis and wound healing (Thiery 2003; Nieto et al. 2016). A mounting body of evidence suggests that EMT is aberrantly activated in cancer cells to mediate tumor recurrence and metastasis (Yang and Weinberg 2008; Thiery et al. 2009). Study of the molecular mechanism of EMT has been largely restricted to cellular signaling and transcriptional regulation. Recently, work from our group has demonstrated that alternative splicing of the gene *CD44* causally contributes to EMT and breast cancer metastasis (Brown et al. 2011; Reinke et al. 2012; Xu et al. 2014; Zhao et al. 2016). Further evidence has also emerged to show the essential role of alternative splicing of other genes in controlling EMT (Lu et al. 2013; Hernandez et al. 2015). A variety of splicing regulatory proteins have also been implicated in EMT alternative splicing, however few have been shown to have essential functional roles during EMT (Warzecha et al. 2010; Braeutigam et al. 2013; Xu et al. 2014; Yang et al. 2016).

Investigating the mechanisms underlying the regulation of *CD44* alternative splicing led us to identify antagonistic roles between two splicing factors, hnRNPM and ESRP1. hnRNPM promotes *CD44* variable exon skipping and favors a mesenchymal phenotype, whereas ESRP1 stimulates *CD44* variable exon inclusion and promotes an epithelial cellular state (Warzecha et al. 2009; Brown et al. 2011; Reinke et al. 2012; Xu et al. 2014). Interestingly, hnRNPM is ubiquitously expressed but functions in a mesenchymal cell-state specific manner to regulate *CD44* alternative splicing. This cell-state restricted activity of hnRNPM is guided in part by competition between ESRP1 and hnRNPM (Xu et al. 2014). hnRNPM and ESRP1 share common guanine-uridine-rich (GU-rich) binding sites (Dittmar et al. 2012; Huelga et al. 2012) and the presence of ESRP1 suppresses the activity of hnRNPM by binding at the same cis-elements in *CD44* introns (Xu et al. 2014). Given the functional consequences of hnRNPM and ESRP1 in modulating EMT, we hypothesized that these two splicing factors compete to regulate not only *CD44* alternative splicing, but also many other splicing events which may be associated with EMT. The balance between hnRNPM and ESRP1 splicing regulation may therefore control the phenotypic switch between an epithelial state and a mesenchymal state.

In this study, we analyzed splicing events regulated by both hnRNPM and ESRP1. Our results show that hnRNPM and ESRP1 exhibit inverse activities in regulating most co-regulated splicing events. Unexpectedly, they also display concordant activities when regulating a subset of splicing events. Importantly, our results reveal that hnRNPM and ESRP1 regulate a set of cassette exons to promote and inhibit EMT, respectively. Discordantly regulated cassette exons are enriched in guanine-uridine (GU) rich motifs specifically in the downstream intron, corresponding with known hnRNPM and ESRP1 binding motifs and likely sites of competitive splicing regulation. This mode of regulation is more widespread than previously appreciated. Co-regulated cassette exons also stratify breast cancer patients by overall survival and correlate with cancer-relevant gene sets, highlighting the importance of hnRNPM and ESRP1 splicing regulation in cancer biology.

## RESULTS AND DISCUSSION

### Splicing factors ESRP1 and hnRNPM co-regulate a set of cassette exons

In an effort to better understand how the key splicing regulators ESRP1 and hnRNPM functionally interact with each other, we compared RNA sequencing datasets in response to hnRNPM and ESRP1 perturbation. We focused on cassette exons, the most common type of alternative splicing (Wang et al. 2008). Alternative splicing levels were quantified using the Percent Spliced In (PSI) metric, which is a measure of the relative abundance of the exon inclusion isoform. We performed RNA sequencing analysis after hnRNPM knockdown in the well-established MDA-MB-231-derived lung and bone metastatic 4175 (LM2) and 1833 (BM1) cell lines (Kang et al. 2003; Minn et al. 2005) and obtained a set of 1635 hnRNPM-regulated alternative cassette exons in these mesenchymal-type cells. Using previously published ESRP1 overexpression and knockdown datasets (Warzecha et al. 2010; Dittmar et al. 2012; Yang et al. 2016), we derived a corresponding set of 1300 ESRP1-regulated cassettes. Intersecting these datasets resulted in a statistically significant overlap of 213 co-regulated cassette exons (Fig. 1A, Supplemental Table S1, p = 7.9e-115, hypergeometric test). Roughly two-thirds of the co-regulated exons (134/213) were regulated discordantly while the remaining exons (79/213) were regulated concordantly (Fig. 1B). The fact that the majority of co-regulated events show discordant regulation mirrors the antagonistic role that ESRP1 and hnRNPM play in favoring cell-state specific splicing programs (Xu et al. 2014). By contrast, the concordant splicing regulation by ESRP1 and hnRNPM suggests that they cooperate to control splicing within a subset of genes. Thus, the co-regulation of splicing between ESRP1 and hnRNPM is not purely antagonistic and may be more complex than previously understood.

**Figure 1:**
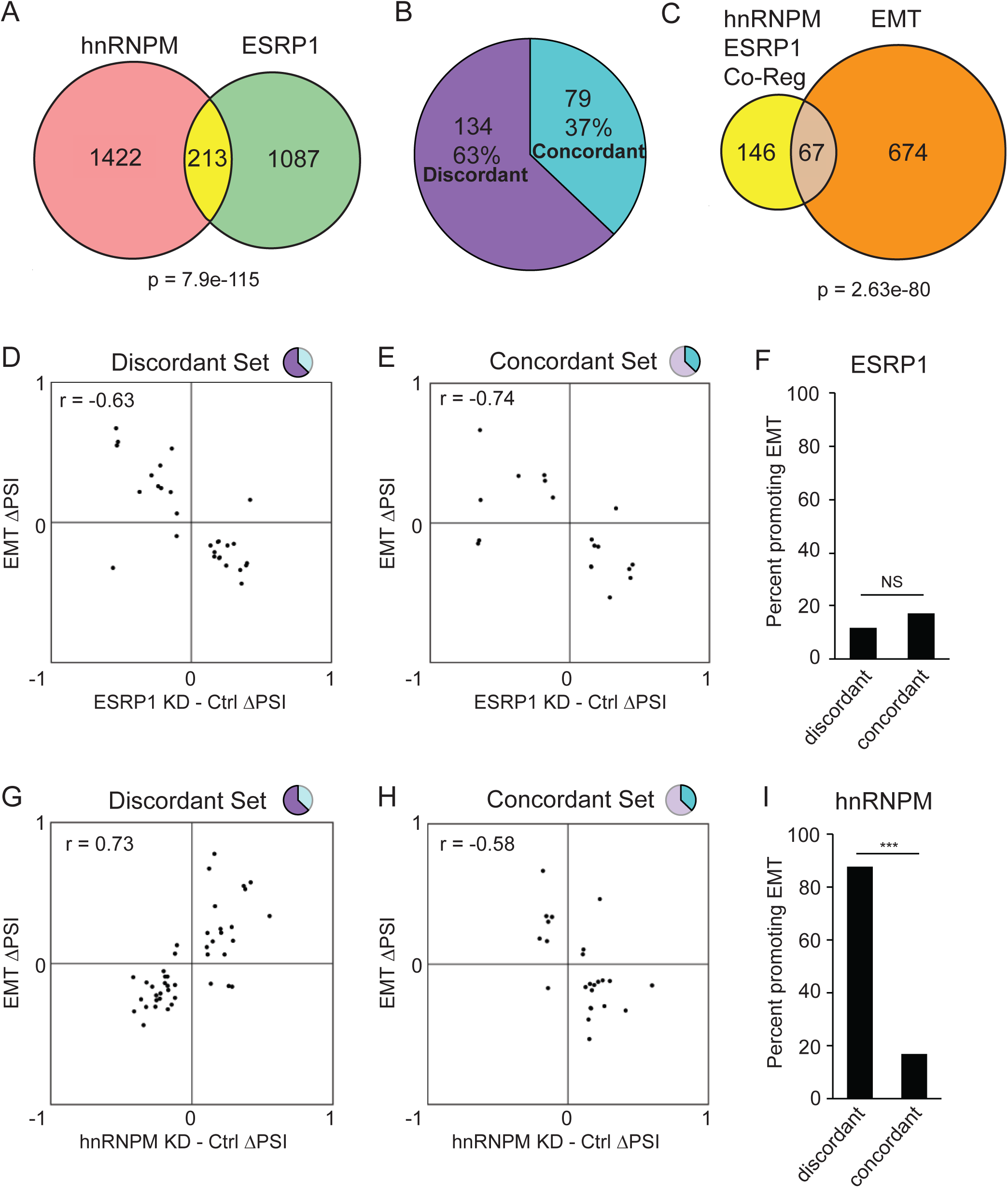
hnRNPM and ESRP1 co-regulate a set of cassette exons in discordant and concordant manners. (A) RNA-sequencing analysis of hnRNPM knockdown and ESRP1 overexpression/knockdown datasets identified splicing factor dependent cassette exon events. In total, 213 cassette exon events showed overlapping regulation by hnRNPM and ESRP1. p-value by hypergeometric test. (B)The majority, 63% (134/213) cassette exons, were regulated discordantly by hnRNPM and ESRP1. The minority, 37% (79/213) cassette exons, were regulated concordantly. (C) 67 of the hnRNPM-ESRP1 co-regulated exons overlap with exons regulated during EMT. p-value by hypergeometric test. (D,E). Both ESRP1 discordantly regulated exons (D) and concordantly regulated exons (E) show a negative correlation with EMT splicing. (F) 12% of ESRP1-regulated discordant exons promote EMT compared to 17% of concordant exons. (G) hnRNPM discordantly regulated exons show a positive correlation with EMT. (H) hnRNPM concordant exons show a negative correlation with EMT. (I). 88% of hnRNPM discordant exons promote EMT compared to 17% of hnRNPM concordant exons. p < 0.001 by Fisher’s exact test.

As hnRNPM and ESRP1 play important roles in regulating alternative splicing during EMT, we overlapped the co-regulated exons with a set of EMT regulated alternative splicing events derived from previous studies (Shapiro et al. 2011; Yang et al. 2016). Over 30% (67/213) of the co-regulated exons overlap with EMT, representing a statistically significant overlap (Fig. 1C, p = 2.63e-80, hypergeometric test). We then analyzed the regulatory roles of hnRNPM and ESRP1 on these EMT-associated splicing events. For both discordant and concordant exons co-regulated by ESRP1 and hnRNPM, ESRP1-mediated splicing inversely correlated with the EMT-associated splicing, in line with the role of ESRP1 as an epithelial specific splicing regulator (Fig. 1D-F) (Warzecha et al. 2010). Interestingly however, hnRNPM showed bi-directional correlation. For the hnRNPM/ESRP1 discordant exons, we observed a positive correlation between hnRNPM-mediated splicing and EMT-associated splicing, indicating that hnRNPM promotes splicing that occurs during EMT (Fig. 1G). For the hnRNPM/ESRP1 concordant exons, however, hnRNPM’s activity inversely correlated with EMT-associated splicing (Fig. 1H). This bi-directional difference in EMT-splicing regulation was statistically significant (Fig. 1I). These results suggest that although the majority of hnRNPM-regulated events are consistent with its role in driving a mesenchymal splicing program in opposition to ESRP1, hnRNPM may also be involved in a small subset of splicing events regulated in favor of an epithelial splicing pattern when functioning in concert with ESRP1.

### Validation of ESRP1 and hnRNPM co-regulated cassette exons

We experimentally validated four of the concordant and four of the discordant co-regulated splicing events using RT-PCR upon shRNA-mediated hnRNPM or ESRP1 knockdown to confirm splicing regulation observed in the RNA sequencing studies (Fig. 2A-C). Given the role of hnRNPM in promoting EMT, we conducted hnRNPM knockdown in the mesenchymal LM2 cells. Because ESRP1 is only expressed in epithelial cells, we performed ESRP1 knockdown in immortalized human mammary epithelial cells (HMLE) (Yang and Weinberg 2008; Xu et al. 2014). The validation set confirms that hnRNPM and ESRP1 co-regulate splicing events in both concordant and discordant manners (Fig. 2C). Moreover, we examined the specificity of the hnRNPM and ESRP1 splicing regulatory relationships by using two splicing minigenes: one contains a discordantly regulated exon at *CD44* variable exon 5 and the other contains a concordant exon at *MARK3* exon 17. Co-transfection experiments of the *CD44v5* minigene with hnRNPM or ESRP1 in 293FT cells showed that hnRNPM promotes v5 exon skipping, whereas ESRP1 inhibits it (Fig. 2D), supporting discordant regulation of hnRNPM and ESRP1. By contrast, co-transfection of the concordant *MARK3* exon 17 minigene with hnRNPM or ESRP1 both resulted in dose-dependent increases in exon skipping, mirroring the concordant regulation of splicing observed in the RNA-sequencing data (Fig. 2E). These results show that hnRNPM and ESRP1 function in both discordant and concordant fashions that are dependent on splicing substrates.

**Figure 2:**
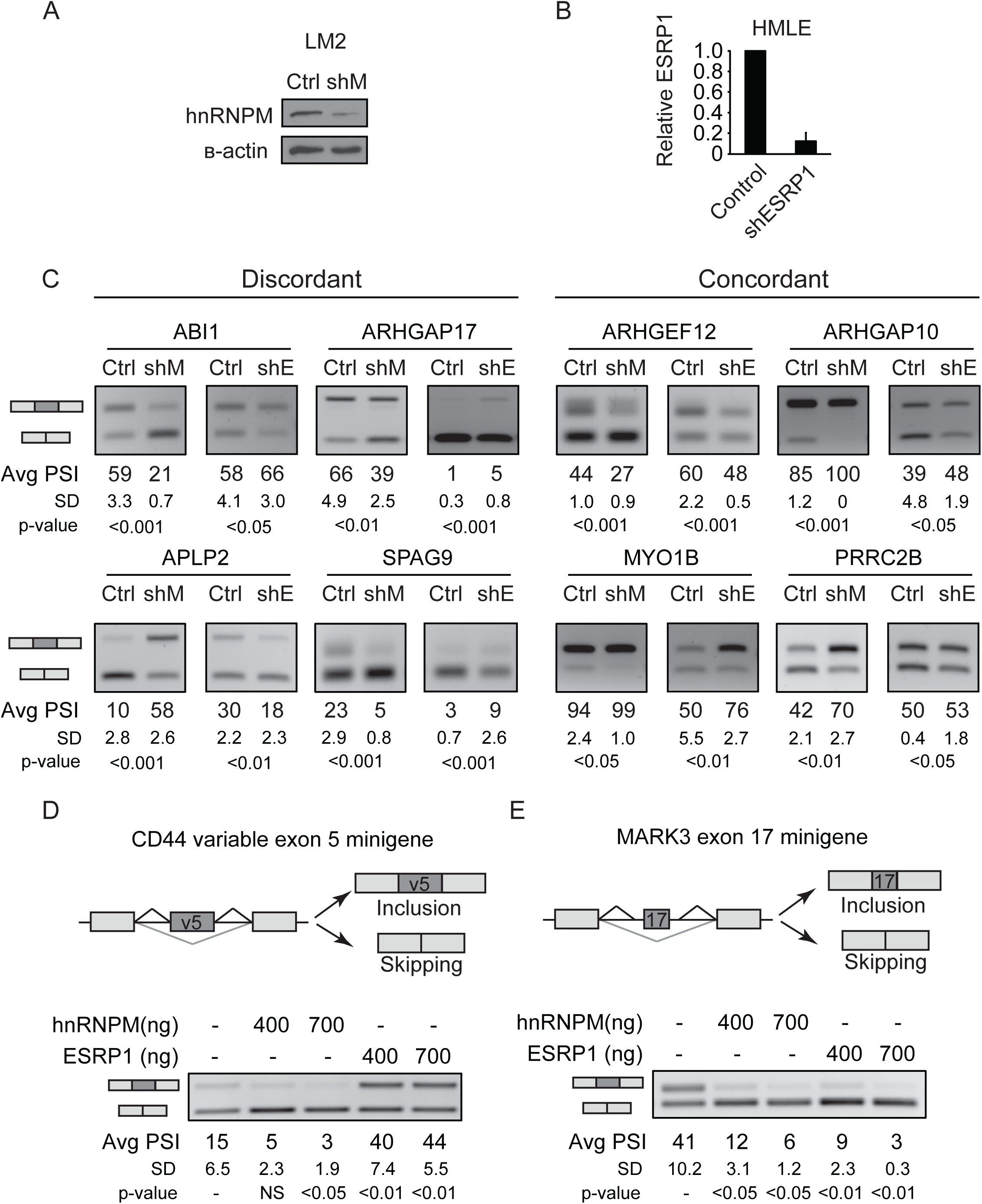
Validation of hnRNPM and ESRP1 co-regulation of cassette exons. (A) hnRNPM knockdown in LM2 cells. (B) ESRP1 knockdown in HMLE cells. (Error bars = S.E.M, n=3) (C) (Left) Cassette exon splicing events regulated discordantly by hnRNPM and ESRP1. (Right) Splicing events regulated concordantly by hnRNPM and ESRP1. Avg PSI represents an average of three experiments. SD represents standard deviation of PSI values. P-value calculated by Student’s T-test comparing KD to control. (D) Splicing minigene analysis of co-transfection of *CD44* variable exon 5 minigene with hnRNPM in 293FT cells promotes exon skipping while ESRP1 promotes exon inclusion. (E) Co-transfection of *MARK3* exon 17 minigene and hnRNPM or ESRP1 in 293FT cells shows that both promote exon skipping. Avg PSI represents an average of three experiments. SD represents standard deviation of PSI values. p-value calculated by Student’s T-test comparing each transfection and control.

### ESRP1 and hnRNPM discordantly regulated exons are enriched in shared binding sites

In order to better understand the functional relationship between hnRNPM and ESRP1 in co-regulating alternative splicing, we performed motif enrichment analysis on the introns near all hnRNPM and ESRP1 co-regulated splicing events (Fig. 3A). Both hnRNPM and ESRP1 are known to bind GU-rich cis-elements primarily in introns (Dittmar et al. 2012; Huelga et al. 2012; Bebee et al. 2015; Yang et al. 2016). We observed selective enrichment of GU-rich hexamers downstream of hnRNPM-repressed and ESRP1-enhanced events, with both sets of events showing the most significant enrichment of the same GUGGUG motif (Fig 3A-C). The observation that GU-motif enrichment was observed downstream of exons regulated oppositely by hnRNPM and ESRP1 suggests that GU-motifs are enriched in discordantly regulated exons. These data support a model where hnRNPM and ESRP1 compete for shared binding sites directly downstream of cassette exons to regulate alternative splicing antagonistically. In addition, we noted significant enrichment of a UGCAUG motif downstream of hnRNPM-enhanced and ESRP1-repressed events. This sequence corresponds to the well-known binding motif of the Rbfox family of RNA binding proteins, of which RBFOX2 has been shown to regulate alternative splicing during EMT (Venables et al. 2013).

**Figure 3:**
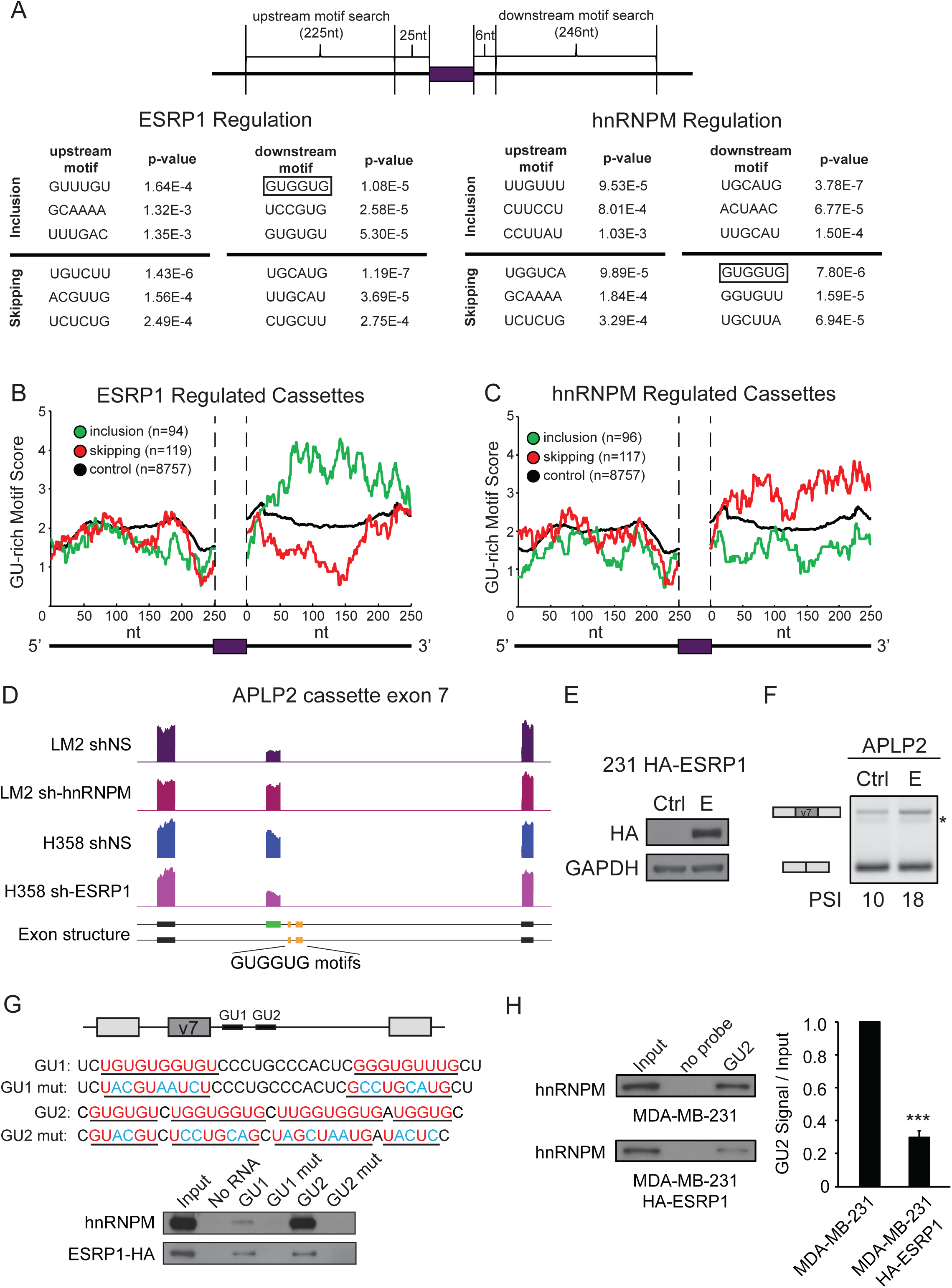
hnRNPM and ESRP1 show common motif enrichment downstream of discordantly regulated exons. (A) K-mer enrichment analysis showing top three enriched 6-mers in introns flanking the 213 hnRNPM and ESRP1 co-regulated cassette exons reveals enrichment of the same top GU-rich motif (GUGGUG, black box) downstream of cassette exons in ESRP1-enhanced and hnRNPM-repressed events. (B,C) RNA motif map analysis of GU-rich motifs in introns flanking hnRNPM/ESRP1 co-regulated exons with respect to ESRP1 regulation (B) and hnRNPM regulation (C) reveals enrichment of GU-rich motifs in downstream introns of ESRP1-enhanced and hnRNPM-repressed events. Inclusion events (green). Skipping events (red). Control events (black). (D) Genome browser plot of RNA sequencing datasets showing hnRNPM and ESRP1 discordantly regulated cassette exon at *APLP2* exon 7. hnRNPM promotes exon skipping while ESRP1 promotes exon inclusion. Black bars indicate constitutive exons. Green bar indicates variable exon 7. Yellow bars indicate location of the top enriched GUGGUG motif within 250 nucleotides downstream of *APLP2* exon 7. (E) Immunoblot of HA-tagged ESRP1 overexpression in MDA-MB-231 cells. (F) ESRP1 overexpression in MDA-MB-231 cells results in increased PSI and exon inclusion of *APLP2* exon 7. * indicates non-specific band. (G) (Upper panel) Two GU-rich regions were identified within 250 nucleotides downstream of *APLP2* exon 7. RNA probes GU1 and GU2 containing stretches of GU nucleotides underlined and in red with mutant probes GU1-mut and GU2-mut with mutated sequences colored in blue. (Lower panel) RNA pull down analysis using RNA probes blotting for endogenous hnRNPM and overexpressed HA-ESRP1 in MDA-MB-231 cells shows hnRNPM and ESRP1 bind common GU-rich sequences. (H) RNA pull down experiments using a static amount of the biotinylated GU2 RNA probe and cell lysate assaying for hnRNPM in the MDA-MB-231 cell line with low ESRP1 expression and the same line with HA-ESRP1 overexpression. 2.5% input was provided as a loading control for both samples. Overexpression of ESRP1 leads to less hnRNPM binding, suggesting that ESRP1 competes for the same GU2 binding site. (Error bars = S.E.M, n=3, *** = p-value < 0.001).

To experimentally examine the binding relationships of hnRNPM and ESRP1, we analyzed their ability to bind to the GU-rich motifs downstream of the discordantly regulated *APLP2* cassette exon 7, which was validated in Fig. 2C. *APLP2* contains three occurrences of the most highly enriched GU-rich motif GUGGUG within 250 nt downstream of *APLP2* cassette exon 7 (Fig. 3D). Experiments were conducted in MDA-MB-231 cells, which do not express ESRP1, and MDA-MB-231 cells ectopically expressing HA-tagged ESRP1 (Fig. 3E). As predicted from the RNA-seq data and validation experiments, an increase in APLP2 PSI was observed upon ESRP1 overexpression (Fig. 3F). In order to examine binding of hnRNPM and ESRP1 to the GU-rich motifs downstream of *APLP2* exon 7, we designed two 5’ biotinylated RNA probes: GU1 and GU2. The GU1-probe encompasses the first GUGGUG motif and a stretch of GU-rich sequences. The GU2-probe contains the other two GUGGUG motifs and additional GU-rich sequences (Fig. 3G, top panel). RNA pull down assays show that both hnRNPM and ESRP1 bind these RNA probes, however hnRNPM shows stronger binding affinity to isomolar concentrations of the GU2 probe compared to the GU1 probe while ESRP1 binds to both probes equally (Fig. 3G, bottom panel). These binding activities are specific because binding of both proteins was abolished upon disruption of GU-rich motifs. To determine whether hnRNPM binding is decreased in the presence of ESRP1, we compared relative hnRNPM binding to the GU2 probe in parental MDA-MB-231 cells compared to MDA-MB-231 cells overexpressing HA-ESRP1. By comparing the amount of hnRNPM that was associated with the GU2 probe relative to input, we found a 70% reduction in hnRNPM binding in the HA-ESRP1-expressing MDA-MB-231 cells (Fig. 3H), suggesting that ESRP1 is capable of competing with hnRNPM for the same binding sites on the APLP2 pre-mRNA.

### ESRP1 and hnRNPM co-regulated exons are enriched in EMT processes and correlate with breast cancer signatures and patient survival

In order to better understand the relevance of the complex regulation of splicing by hnRNPM and ESRP1 to disease phenotypes, we performed gene ontology analysis using DAVID on the set of 213 co-regulated exons (Huang da et al. 2009b; Huang da et al. 2009a). We observed significant GOTERMs associated with cell polarity, cell adhesion, and cytoskeletal dynamics (Fig. 4A), all processes that are critical for EMT (Supplemental Table S2). To assess the contribution of hnRNPM and ESRP1 co-regulation of splicing in breast cancer patient samples, we mined the publicly available The Cancer Genome Atlas (TCGA) RNA-sequencing data for invasive breast carcinoma (BRCA) and calculated the PSI values for hnRNPM and ESRP1 co-regulated splicing events (Cancer Genome Atlas 2012). After stratifying the patients based on PSI levels for each exon via 2-means clustering, we observed significant differences in patient survival based on the splicing levels of 13 cassette exons (FDR < 0.05, log rank test). As an example, *SPAG9* exon 24 inclusion is increased during EMT and this exon inclusion is stimulated by hnRNPM and inhibited by ESRP1. We found that *SPAG9* exon 24 inclusion levels predict poorer patient survival (Fig. 4B). In addition, *ZMYND8* exon 22 undergoes exon skipping during EMT and the exon skipping event is promoted by hnRNPM and antagonized by ESRP1. TCGA data analysis showed that *ZMYND8* exon inclusion is associated with a better prognosis in breast cancer (Fig. 4B).

**Figure 4:**
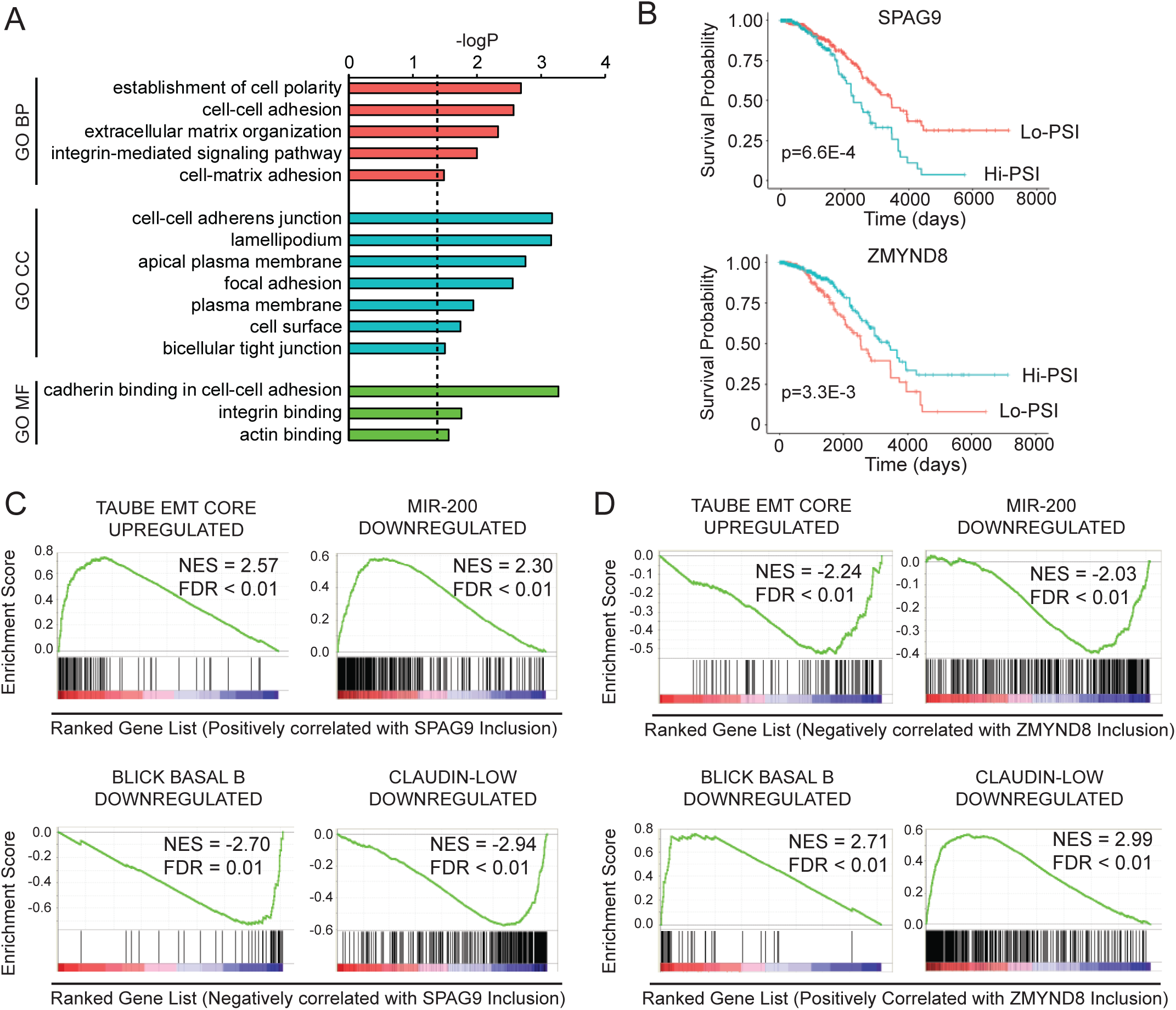
hnRNPM-ESRP1 co-regulated exons are associated with EMT and breast cancer survival. (A) Gene ontology analysis of 213 hnRNPM-ESRP1 co-regulated exons representing 184 genes contain significant terms associated with cell polarity, adhesion, migration, and the cytoskeleton. Direct GOTERMS BP, MF, and CC were queried using DAVID. (B) hnRNPM/ESRP1 discordantly regulated exon inclusion levels, such those of SPAG9 and ZMYND8, stratify breast cancers patients by overall survival. (C) Genes positively correlated with SPAG9 inclusion are upregulated during EMT and downregulated by mir-200 while genes negatively correlated with SPAG9 inclusion are downregulated in basal and claudin-low breast cancer subtypes. (D) Conversely, genes upregulated during EMT and downregulated by mir-200 are negatively correlated with ZMYND8 inclusion while genes downregulated in basal and claudin-low breast cancers are positively correlated with ZMYND8 inclusion.

Intrigued by the role these exons played in predicting breast-cancer patient survival, we performed Gene Set Enrichment Analysis (GSEA) on gene signatures that are correlated with *SPAG9* and *ZMYND8* inclusion. Within the TCGA dataset, a published EMT gene signature was positively correlated with *SPAG9* exon inclusion, while this signature was negatively correlated with *ZMYND8* inclusion (Taube et al. 2010) (Fig. 4C,D). Genes downregulated by mir-200, a potent repressor of EMT, also showed a positive correlation with *SPAG9* exon inclusion but a negative correlation with *ZMYND8* inclusion (Korpal and Kang 2008; Park et al. 2008). Moreover, genes downregulated in basal subtype and claudin-low subtype breast cancers, which are generally non-hormone dependent and resistant to conventional therapies, were negatively correlated with *SPAG9* exon inclusion (Blick et al. 2010; Prat et al. 2010). Conversely, these genes were positively correlated with *ZMYND8* inclusion.

### Conclusions

In summary, this study reveals widespread co-regulation of alternative splicing by hnRNPM and ESRP1, both identified as key regulators of EMT splicing programs (Warzecha et al. 2010; Xu et al. 2014). hnRNPM-regulated cassette exons significantly overlap with ESRP1-regulated cassette exons, with the majority of co-regulated events showing discordant splicing regulation, suggesting that hnRNPM and ESRP1 largely serve to functionally antagonize one another. We also observed a subset of splicing events hnRNPM and ESRP1 concordantly regulate that inversely correlate with EMT splicing. Although these events represent a minority of co-regulated events, these results suggest that hnRNPM is partly correlated with antagonistic regulation of splicing during EMT. These results are surprising as we observed that hnRNPM is required for cells to undergo EMT (Xu et al. 2014). Interestingly, this scenario is reminiscent of that observed for RBM47, an RNA binding protein that inhibits EMT (Vanharanta et al. 2014). RBM47 showed primarily discordant regulation of splicing events compared to EMT, but also concordant regulation of a subset of splicing events that promote EMT (Yang et al. 2016). These findings highlight the importance of understanding the combinatorial regulation of splicing by different factors with respect to a complex biological process such as EMT. Whether the hnRNPM-regulated splicing events that oppose EMT play a functional role during EMT or are important for regulating cellular processes that are highly active in epithelial cell states will be an interesting area for future study.

Some of the alternative splicing events co-regulated by hnRNPM and ESRP1 have been investigated in detail to understand their functional contributions to EMT. *CD44* contains a set of variable exons that undergo extensive alternative splicing during EMT. Exon skipping of all variable exons of *CD44* to generate the CD44s isoform is required for EMT and has been shown to promote Akt-signaling, mediate invadopodia activity, and attenuate degradation of EGFR to promote sustained RTK signaling, all of which have implications during EMT (Brown et al. 2011; Zhao et al. 2016; Liu and Cheng 2017; Wang et al. 2017). ESRP1 and hnRNPM directly regulate alternative splicing of *CD44* in a discordant manner, with ESRP1 driving exon inclusion and hnRNPM promoting exon skipping. Another hnRNPM/ESRP1 co-regulated splicing event that plays a role during EMT is alternative splicing of *EXOC7.* The EXOC7 mesenchymal isoform is capable of promoting actin polymerization and increased cell invasion compared to the epithelial isoform (Lu et al. 2013). ESRP1 was shown to regulate *EXOC7* splicing to promote production of the epithelial isoform, although the role of hnRNPM is not known. In addition, hnRNPM and ESRP1 co-regulate exon skipping of *TCF7L2.* The exon skipping *TCF7L2* isoform is capable of greater activation of Wnt signaling cascade target genes (Weise et al. 2010), and this event is upregulated in the mesenchymal state. It is worth noting that in our study all three of these splicing events are discordantly regulated by hnRNPM and ESRP1, with ESRP1 promoting production of the epithelial isoform and hnRNPM promoting the mesenchymal splicing pattern. While the precise molecular consequences of most of the hnRNPM/ESRP1 regulated splicing events during EMT are unknown, our observation that hnRNPM and ESRP1 co-regulated splicing events are enriched in genes associated with cytoskeleton remodeling and cell adhesion present interesting avenues for further study.

Our study on motif analysis suggests a common mode of competitive interaction at wide-spread discordantly regulated exons where hnRNPM and ESRP1 compete for binding sites. In addition to enrichment of GU-rich motifs, we also observed striking enrichment of the UGCAUG motif downstream of hnRNPM-enhanced and ESRP1-repressed events. This UGCAUG motif corresponds to the highly conserved Rbfox family of RNA binding proteins.

Rbfox motifs have been shown previously to enrich near exons regulated by hnRNPM (Damianov et al. 2016) and ESRP1 (Warzecha et al. 2010; Yang et al. 2016). RBFOX2 was recently shown to function in a complex with hnRNPM along with other splicing factors (Damianov et al. 2016). Since RBFOX2 has been shown to promote alternative splicing during EMT (Shapiro et al. 2011; Venables et al. 2013), we speculate that RBFOX2 may function in concert with hnRNPM and ESRP1 to regulate EMT-associated alternative splicing. In-depth analysis of hnRNPM and ESRP1 splicing regulation informs our understanding of splicing factor binding and functional dynamics in the context of disease-relevant splicing programs and indicates the importance of understanding the competitive and cooperative mechanisms of splicing regulation that allow precise modulation of alternative splicing.

Taken together, we show that co-regulation of alternative splicing by hnRNPM and ESRP1 is wide-spread and primarily antagonistic, although a subset of events is regulated concordantly. Furthermore, we demonstrate that in controlling hnRNPM/ESRP1 discordantly regulated events, hnRNPM promotes alternative splicing in the same direction as EMT. hnRNPM and ESRP1 splicing antagonism is explained by competition for GU-rich elements downstream of co-regulated exons. Lastly, hnRNPM/ESRP1 co-regulated splicing correlates with EMT and breast cancer-associated gene sets and predicts breast cancer patient survival. Taken together, this study highlights the complex regulation of alternative splicing by ESRP1 and hnRNPM as well as the relevance of this regulatory interaction in EMT and cancer.

## MATERIALS AND METHODS

### Cell Lines

Maintenance of immortalized human mammary epithelial cells (HMLE) cells was conducted as previously described (Mani et al. 2008). Human embryonic kidney 293FT, human breast carcinoma MDA-MB-231, and MDA-MB-231 metastatic derivative lines 4175 (LM2) and 1833 (BM1) were grown in DMEM supplemented with 10% FBS, L-glutamine, penicillin, and streptomycin. The HA-ESRP1 overexpressing MDA-MB-231 cell line was described previously (Reinke et al. 2012).

### Plasmids, shRNAs, and ESRP1 overexpression

Two expression plasmids, pcDNA3-FLAG-hnRNPM and pcDNA3-FLAG-ESRP1 were subcloned from pECFP-hnRNPM (Lleres et al. 2010) and pBRIT-ESRP1 (Brown et al. 2011), respectively. The *CD44* variable exon 5 minigene was described previously (Brown et al. 2011). The *MARK3* exon 17 minigene was constructed through PCR amplification of *MARK3* exon 17 and approximately 400 nucleotides of flanking intron followed by cloning into the BamH1 site of the *CD44v5* minigene. Primers for *MARK3* minigene construction are listed in Supplemental Table S3. The control, hnRNPM, and ESRP1 shRNAs were described previously (Brown et al. 2011; Xu et al. 2014). sh-control = 5’-CCCGAATTAGC TGGACACTCAA-3’. sh-hnRNPM = 5’-GGAATGGAAGGCATAGGATTT-3’. The ESRP1 shRNA was obtained from Open Biosystems (clone V2LHS_155253).

### Transfection, semi-quantitative RT-PCR, and qRT-PCR

Briefly, 2.25 × 10^5^ HEK293FT cells were plated in 24-well plates twenty-four hours prior to transfection. Co-transfection of hnRNPM and/or ESRP1 with splicing minigenes was performed using Lipofectamine 2000 (Invitrogen) per the manufacturer’s instructions. For HEK293FT transfections, 100 ng of minigene was used. RNA was extracted from cells using the E.Z.N.A. Total RNA Kit (Omega Bio-Tek). RNA concentration and purity was measured using a Nanodrop 2000 (Thermo Fisher Scientific). cDNA was generated via reverse transcription using the GoScript Reverse Transcription System (Promega) with 1 μL GoScript RT and 250 ng of RNA in a total volume of 20 μL followed by incubation at 25°C for 5 mins, 42°C for 30 mins, and 70°C for 15 min. Semi-quantitative RT-PCR assaying for splicing products was performed using Hot StarTaq DNA polymerase (Qiagen), and PCR cycles were run for 30 or fewer cycles.

Primers for semi-quantitative analysis were designed in constitutive exons flanking each variable exon (Harvey and Cheng 2016). Semi-quantitative PCR generates both exon inclusion and skipping products which were separated through agarose gel electrophoresis. PCR product intensity was measured using ImageJ image analysis software. qRT-PCR was performed in a total volume of 20 μL with 12.5 ng cDNA and 0.75 μM primers using GoTaq qPCR Master Mix (Promega) per the manufacturer’s instructions on a CFX Connect Real-Time PCR system (BioRad) using a three-step protocol and supplied software. For every qRT-PCR sample per biological replicate, two technical replicates were performed and Ct counts were averaged. Normalization and quantification of qRT-PCR data was done using the 2^-ΔΔCt^ method (Livak and Schmittgen 2001). Primer specificity was checked with melt-curve analysis during qRT-PCR. Primers for semi-quantitative and qRT-PCR analysis were designed to generate products spanning introns (Supplemental Table S3).

### Quantitative immunoblotting

Whole cell lysates or RNA pull down samples were separated by 10% SDS-PAGE, transferred to a PVDF membrane (BioRad), and probed with the appropriate antibody. Primary antibodies used in western blots included HA-HRP 1:1000 (Roche Applied Science), hnRNPM 1:100000 (OriGene), GAPDH (GE) and β-actin (Sigma-Aldrich) were used as a loading controls. After incubation with HRP-tagged secondary antibodies, if appropriate, blots were visualized via chemiluminescence (Thermo-Fisher).

### RNA Pulldown Assays

5’-biotinylated nucleotides were used for RNA pull down experiments. The *APLP2* exon 7 associated probes include GU1: 5’-biotin-UCUGUGUGGUGUCCCUGCCCACUCGGGUGUUUGCU, which was mutated to GU1 mut: 5’-biotin-UCUACGUAAUCUCCCUGCCCACUCGCCUGCAUGCU and GU2: 5’-biotin-CGUGUGUCUGGUGGUGCUUGGUGGUGAUGGUGC, which was mutated to GU2 mut: 5’-biotin-CGUACGUCUCCUGCAGCUAGCUAAUGAUACUCC. Biotinylated RNA oligos (10 μL at 40 μM) were immobilized on 50 μL of streptavidin beads (50% slurry; Thermo Fisher) in a total volume of 400 μL 1X binding buffer (20 mM Tris, 200 mM NaCl, 6 mM EDTA, 5 mM sodium fluoride, 5 mM β-glycerophosphate, 2 mg/mL aprotinin, pH 7.5) for 2 hours at 4°C in a rotating shaker. After immobilization, beads were washed 3 times in 1X binding buffer, then 200 μg MDA-MB-231 or MDA-MB-231 HA-ESRP1 cell lysates were suspended with the beads in 400 μL of 1X binding buffer for incubation at 4°C overnight. Beads were then washed 3 times in 1X binding buffer, resuspended in 60 μL of 2X Laemmli sample buffer (Bio-Rad), and boiled for 5 m. Ten μL of sample was analyzed under denaturing conditions on 10% SDS-PAGE and detected via immunoblotting.

### RNA Sequencing Analysis

RNA was extracted from LM2 or BM1 cells stably expressing sh-control or sh-hnRNPM using Trizol, and poly-A-selected RNA-seq libraries were generated using TruSeq stranded mRNA library preparation kits (Illumina) and subjected to 100-base-pair PE stranded RNA-seq on an Illumina HiSeq 4000. RNA-seq reads were aligned to the human genome (GRCh37, primary assembly) and transcriptome (Gencode version 24 backmap 37 comprehensive gene annotation) using STAR version 2.5.3a (Dobin et al. 2013) using the following parameters: STAR ‐‐runThreadN 16 - -alignEndsType EndToEnd ‐‐quantMode GeneCounts ‐‐outSAMtype BAM SortedByCoordinate. Differential alternative splicing was quantified using rMATS version 3.2.5 (Shen et al. 2014) using the following nondefault parameters: -t paired -len 100 -analysis U -libType fr-firststrand. and the following cutoffs: FDR < 0.05, ΔPSI ≥ 0.1, and average junction reads per cassette event ≥10. Control cassette exons were identified by the following filters: FDR > 0.5, minimum PSI for sh-control or sh-hnRNPM <0.85, maximum PSI > 0.15, and average junction reads per cassette event per replicate ≥10. These filters were selected to identify cassette exons with evidence of alternative splicing but were not differentially spliced upon hnRNPM depletion. RNA sequencing data for ESRP1 was processed in the same way after retrieving data from GEO record GSE74592. Other ESRP1-regulated cassette exons were obtained from the supplemental materials of published studies (Warzecha et al. 2010; Dittmar et al. 2012). EMT-regulated cassette exons were obtained from the supplemental materials of two published studies (Shapiro et al. 2011; Yang et al. 2016). Sequencing datasets for hnRNPM are deposited in the Gene Expression Omnibus at GSE112516.

### Motif Enrichment and RNA Motif Maps

Kmer enrichments were calculated using 250 bp of the sequence flanking the cassette exons. To avoid enrichment of splice site motifs, we removed 9 nucleotides downstream of 5’ splice sites and 25 nucleotides upstream of 3’ splice sites. To assess enrichment of hexamers, we adapted a previously published method in a custom python script (Coelho et al. 2015). Given a set of test sequences and control sequences, the frequency of occurrence of each possible hexamer of RNA (4096) was computed per nucleotide in the control sequences to establish a background frequency. Then for each hexamer for each sequence in the test or control set, the frequency of the hexamer per nucleotide was calculated and compared to the established background frequency for that hexamer. To determine if the hexamer was enriched in the test set relative to the control set, a one-tailed Fisher’s exact test was conducted using the sum of sequences with a hexamer frequency greater than background and the sum of sequences with a hexamer frequency less than background for the test set and control set of sequences.

For the GU-rich motif RNA map analysis, the top 12 6-mer motifs from an ESRP1-SELEX-Seq analysis were obtained (Dittmar et al. 2012). The motifs were UGGUGG, GGUGGG, GUGGUG, GUGGGG, GUGUGG, GGUGUG, UGUGGG, GGUGGU, GUGGGU, UGGGGU, GGGGGU, UGGGGG). The 250 bp of sequence flanking upstream and downstream hnRNPM and ESRP1 regulated cassette exons as well as control exons obtained from the rMATS alternative splicing analysis were also obtained. The GU-rich motif score was computed in a custom python script by counting the number of nucleotides covered by any of the GU-rich motifs in a sliding window of 50 bp shifted 1-nucleotide at a time across the 250 bp interval in all of the regulated and control cassette exons. The GU-rich motif score was set equal to the percent of nucleotides covered by the motifs in each of the sliding windows and plotted for regulated exons, stratified by inclusion or skipping, and control exons.

### TCGA BRCA Survival Analysis, GSEA, and Gene Ontology

Processed TCGA BRCA level 3 RNA Seq V2 data for exon junctions and gene expression were downloaded from the Genomic Data Commons Legacy Archive (Cancer Genome Atlas 2012). Cassette exon PSI values in each patient sample were calculated using the following equation from the exon junction files: PSI = (Inclusion junction reads / 2) / ((Inclusion junction reads / 2) + (Skipping junction reads)). Patients were clustered into two groups using K-means clustering. Kaplan-Meier survival analysis was conducted between these two groups and p-values were computed using log-rank tests.

For GSEA analysis, correlation values between the given cassette exon PSI values and all genes in the TCGA BRCA RNA-seq V2 datasets were computed. Genes were then ranked by correlation and GSEA was performed using the Broad Institute javaGSEA desktop application (Mootha et al. 2003; Subramanian et al. 2005).

Gene ontology analysis was conducted using DAVID v6.8 with the gene list composed of all genes containing an exon in the 213 hnRNPM-ESRP1 co-regulated exons (Huang da et al. 2009b; Huang da et al. 2009a). GOTERM enrichment was restricted to GOTERM BP-DIRECT, GOTERM MF-DIRECT, and GOTERM CC-DIRECT. The background gene set was composed of all genes in hnRNPM and ESRP1 RNA sequencing datasets with FPKM > 3 in at least one sample.

### Statistics

Statistical analyses included two-tailed Student’s T-tests and hypergeometric testing unless otherwise noted. *P*-values < 0.05 were considered significant. *P* < 0.05 (*), *P* < 0.01 (**), *P* < 0.001 (***).

## SUPPLEMENTAL MATERIAL

Supplemental material is available for this article.

## ACKNOWLEDGEMENTS

This work was supported in part by the US National Institutes of Health Ruth L. Kirschstein National Research Service Award [1F30CA196118 to S.E.H.] and US National Institutes of Health R01s [CA182467 and GM110146 to C. C., HG006264 and HG009417 to X.X.]. C. C. is a Cancer Prevention Research Institute of Texas Scholar in Cancer Research.

## Authors’ contributions

S.E.H. and C.C. designed experiments. Y.X. generated cell lines. X.L. and X.D.G. cloned splicing minigenes. Y.Q. performed splicing quantification from TCGA data. S.E.H. performed experiments, bioinformatics analysis, and analyzed data. J.A. assisted with analysis of RNA-sequencing data. X.X. supervised bioinformatics analysis. S.E.H. and C.C. wrote the manuscript. C.C. supervised the study.

## REFERENCES

Barash Y, Calarco JA, Gao W, Pan Q, Wang X, Shai O, Blencowe BJ, Frey BJ. 2010. Deciphering the splicing code. Nature 465.

Bebee TW, Park J, Sheridan KI, Warzecha CC, Cieply BW, Rohacek AM, Xing Y, Carstens RP. 2015. The splicing regulators Esrp1 and Esrp2 direct an epithelial splicing program essential for mammalian development. eLife 4.

Blick T, Hugo H, Widodo E, Waltham M, Pinto C, Mani SA, Weinberg RA, Neve RM, Lenburg ME, Thompson EW. 2010. Epithelial mesenchymal transition traits in human breast cancer cell lines parallel the CD44(hi/)CD24 (lo/-) stem cell phenotype in human breast cancer. J Mammary Gland Biol Neoplasia 15: 235–252.

Braeutigam C, Rago L, Rolke A, Waldmeier L, Christofori G, Winter J. 2013. The RNA-binding protein Rbfox2: an essential regulator of EMT-driven alternative splicing and a mediator of cellular invasion. Oncogene 33.

Brown RL, Reinke LM, Damerow MS, Perez D, Chodosh LA, Yang J, Cheng C. 2011. CD44 splice isoform switching in human and mouse epithelium is essential for epithelial-mesenchymal transition and breast cancer progression. Journal of Clinical Investigation 121.

Cancer Genome Atlas N. 2012. Comprehensive molecular portraits of human breast tumours. Nature 490: 61–70.

Cieply B, Carstens RP. 2015. Functional roles of alternative splicing factors in human disease. Wiley Interdiscip Rev RNA 6: 311–326.

Coelho MB, Attig J, Bellora N, König J, Hallegger M, Kayikci M, Eyras E, Ule J, Smith CWJ. 2015. Nuclear matrix protein Matrin3 regulates alternative splicing and forms overlapping regulatory networks with PTB. The EMBO Journal 34: 653–668.

Damianov A, Ying Y, Lin C-H, Lee J-A, Tran D, Vashisht AA, Bahrami-Samani E, Xing Y, Martin KC, Wohlschlegel JA et al. 2016. Rbfox Proteins Regulate Splicing as Part of a Large Multiprotein Complex LASR. Cell 165: 606–619.

Dittmar KA, Jiang P, Park J, Amirikian K, Wan J, Shen S, Xing Y, Carstens RP. 2012. Genome-wide determination of a broad ESRP-regulated posttranscriptional network by high-throughput sequencing. Molecular and Cellular Biology 32.

Dobin A, Davis CA, Schlesinger F, Drenkow J, Zaleski C, Jha S, Batut P, Chaisson M, Gingeras TR. 2013. STAR: ultrafast universal RNA-seq aligner. Bioinformatics (Oxford, England) 29: 15–21.

ENCODE Project Consortium. 2012. An integrated encyclopedia of DNA elements in the human genome. Nature 489.

Harrow J, Frankish A, Gonzalez JM, Tapanari E, Diekhans M, Kokocinski F, Aken BL, Barrell D, Zadissa A, Searle S et al. 2012. GENCODE: the reference human genome annotation for The ENCODE Project. Genome Res 22: 1760–1774.

Harvey SE, Cheng C. 2016. Methods for Characterization of Alternative RNA Splicing BT - Long Non-Coding RNAs. Springer New York 1402.

Hernandez JR, Kim JJ, Verdone JE, Liu X, Torga G, Pienta KJ, Mooney SM. 2015. Alternative CD44 splicing identifies epithelial prostate cancer cells from the mesenchymal counterparts. Medical Oncology 32.

Huang da W, Sherman BT, Lempicki RA. 2009a. Bioinformatics enrichment tools: paths toward the comprehensive functional analysis of large gene lists. Nucleic Acids Res 37: 1–13.

Huang da W, Sherman BT, Lempicki RA. 2009b. Systematic and integrative analysis of large gene lists using DAVID bioinformatics resources. Nat Protoc 4: 44–57.

Huelga SC, Vu AQ, Arnold JD, Liang TY, Liu PP, Yan BY, Donohue J, Shiue L, Hoon S, Brenner S et al. 2012. Integrative genome-wide analysis reveals cooperative regulation of alternative splicing by hnRNP proteins. Cell Reports 1.

Kang Y, Siegel PM, Shu W, Drobnjak M, Kakonen SM, Cordon-Cardo C, Guise TA, Massague J. 2003. A multigenic program mediating breast cancer metastasis to bone. Cancer cell 3: 537–549.

Korpal M, Kang Y. 2008. The emerging role of miR-200 family of microRNAs in epithelial-mesenchymal transition and cancer metastasis. RNA Biol 5: 115–119.

Liu S, Cheng C. 2013. Alternative RNA splicing and cancer. Wiley Interdisciplinary Reviews: RNA 4.

Liu S, Cheng C. 2017. Akt Signaling Is Sustained by a CD44 Splice Isoform–Mediated Positive Feedback Loop. Cancer Research 77: 3791–3801.

Livak KJ, Schmittgen TD. 2001. Analysis of relative gene expression data using real-time quantitative PCR and the 2(T)(-Delta Delta C) method. Methods 25: 402–408.

Lleres D, Denegri M, Biggiogera M, Ajuh P, Lamond AI. 2010. Direct interaction between hnRNP-M and CDC5L/PLRG1 proteins affects alternative splice site choice. EMBO Rep 11: 445–451.

Lu H, Liu J, Liu S, Zeng J, Ding D, Carstens RP, Cong Y, Xu X, Guo W. 2013. Exo70 isoform switching upon epithelial-mesenchymal transition mediates cancer cell invasion. Developmental Cell 27.

Mani SA, Guo W, Liao MJ, Eaton EN, Ayyanan A, Zhou AY, Brooks M, Reinhard F, Zhang CC, Shipitsin M et al. 2008. The epithelial-mesenchymal transition generates cells with properties of stem cells. Cell 133.

Minn AJ, Gupta GP, Siegel PM, Bos PD, Shu W, Giri DD, Viale A, Olshen AB, Gerald WL, Massague J. 2005. Genes that mediate breast cancer metastasis to lung. Nature 436: 518–524.

Mootha VK, Lindgren CM, Eriksson KF, Subramanian A, Sihag S, Lehar J, Puigserver P, Carlsson E, Ridderstrale M, Laurila E et al. 2003. PGC-1alpha-responsive genes involved in oxidative phosphorylation are coordinately downregulated in human diabetes. Nat Genet 34: 267–273.

Nieto AM, Huang R, Jackson RA, Thiery J. 2016. EMT: 2016. Cell 166.

Pan Q, Shai O, Lee LJ, Frey BJ, Blencowe BJ. 2008. Deep surveying of alternative splicing complexity in the human transcriptome by high-throughput sequencing. Nature Genetics 40.

Park SM, Gaur AB, Lengyel E, Peter ME. 2008. The miR-200 family determines the epithelial phenotype of cancer cells by targeting the E-cadherin repressors ZEB1 and ZEB2. Genes & development 22: 894–907.

Prat A, Parker JS, Karginova O, Fan C, Livasy C, Herschkowitz JI, He X, Perou CM. 2010. Phenotypic and molecular characterization of the claudin-low intrinsic subtype of breast cancer. Breast cancer research: BCR 12: R68.

Reinke LM, Xu Y, Cheng C. 2012. Snail represses the splicing regulator epithelial splicing regulatory protein 1 to promote epithelial-mesenchymal transition. Journal of Biological Chemistry 287.

Shapiro IM, Cheng AW, Flytzanis NC, Balsamo M, Condeelis JS, Oktay MH, Burge CB, Gertler FB. 2011. An EMT-driven alternative splicing program occurs in human breast cancer and modulates cellular phenotype. PLoS Genetics 7.

Shen S, Park J, Lu Z-x, Lin L, Henry MD, Wu Y, Zhou Q, Xing Y. 2014. rMATS: robust and flexible detection of differential alternative splicing from replicate RNA-Seq data. Proceedings of the National Academy of Sciences of the United States of America 111: E5593–5601.

Subramanian A, Tamayo P, Mootha VK, Mukherjee S, Ebert BL, Gillette MA, Paulovich A, Pomeroy SL, Golub TR, Lander ES et al. 2005. Gene set enrichment analysis: a knowledge-based approach for interpreting genome-wide expression profiles. Proc Natl Acad Sci U S A 102: 15545–15550.

Taube JH, Herschkowitz JI, Komurov K, Zhou AY, Gupta S, Yang J, Hartwell K, Onder TT, Gupta PB, Evans KW et al. 2010. Core epithelial-to-mesenchymal transition interactome gene-expression signature is associated with claudin-low and metaplastic breast cancer subtypes. Proceedings of the National Academy of Sciences 107.

Thiery J. 2003. Epithelial-mesenchymal transitions in development and pathologies. Current Opinion in Cell Biology 15.

Thiery JP, Acloque H, Huang RY, Nieto MA. 2009. Epithelial-mesenchymal transitions in development and disease. Cell 139.

Vanharanta S, Marney CB, Shu W, Valiente M, Zou Y, Mele A, Darnell RB, Massagué J. 2014. Loss of the multifunctional RNA-binding protein RBM47 as a source of selectable metastatic traits in breast cancer. eLife 3.

Venables JP, Brosseau J-P, Gadea G, Klinck R, Prinos P, Beaulieu J-F, Lapointe E, Durand M, Thibault P, Tremblay K et al. 2013. RBFOX2 Is an Important Regulator of Mesenchymal Tissue-Specific Splicing in both Normal and Cancer Tissues. Molecular and cellular biology 33: 396–405.

Wang ET, Sandberg R, Luo S, Khrebtukova I, Zhang L, Mayr C, Kingsmore SF, Schroth GP, Burge CB. 2008. Alternative isoform regulation in human tissue transcriptomes. Nature 456: 470–476.

Wang W, Zhang H, Liu S, Kim C, Xu Y, Hurley LA, Nishikawa R, Nagane M, Hu B, Stegh AH et al. 2017. Internalized CD44s splice isoform attenuates EGFR degradation by targeting Rab7A. Proceedings of the National Academy of Sciences 114: 8366–8371.

Warzecha CC, Jiang P, Amirikian K, Dittmar KA, Lu H, Shen S, Guo W, Xing Y, Carstens RP. 2010. An ESRP-regulated splicing programme is abrogated during the epithelial-mesenchymal transition. The EMBO Journal 29.

Warzecha CC, Shen S, Xing Y, Carstens RP. 2009. The epithelial splicing factors ESRP1 and ESRP2 positively and negatively regulate diverse types of alternative splicing events. RNA biology 6: 546–562.

Weise A, Bruser K, Elfert S, Wallmen B, Wittel Y, Wohrle S, Hecht A. 2010. Alternative splicing of Tcf7l2 transcripts generates protein variants with differential promoter-binding and transcriptional activation properties at Wnt/beta-catenin targets. Nucleic Acids Res 38: 1964–1981.

Xu Y, Gao XD, Lee JH, Huang H, Tan H, Ahn J, Reinke LM, Peter ME, Feng Y, Gius D et al. 2014. Cell type-restricted activity of hnRNPM promotes breast cancer metastasis via regulating alternative splicing. Genes & Development 28.

Yang J, Weinberg RA. 2008. Epithelial-mesenchymal transition: at the crossroads of development and tumor metastasis. Developmental Cell 14.

Yang Y, Park J, Bebee TW, Warzecha CC, Guo Y, Shang X, Xing Y, Carstens RP. 2016. Determination of a Comprehensive Alternative Splicing Regulatory Network and Combinatorial Regulation by Key Factors during the Epithelial-to-Mesenchymal Transition. Molecular and cellular biology 36.

Zhao P, Xu Y, Wei Y, Qiu Q, Chew T-L, Kang Y, Cheng C. 2016. The CD44s splice isoform is a central mediator for invadopodia activity. J Cell Sci 129.

